# Regulome-wide association study identifies enhancer properties associated with risk for schizophrenia

**DOI:** 10.1101/2021.06.14.448418

**Authors:** Alex M. Casella, Carlo Colantuoni, Seth A. Ament

## Abstract

Genetic risk for complex traits is strongly enriched in non-coding genomic regions involved in gene regulation, especially enhancers. However, we lack adequate tools to connect the characteristics of these disruptions to genetic risk. Here, we propose RWAS (Regulome Wide Association Study), a new framework to identify the characteristics of enhancers that contribute to genetic risk for disease. Applying our technique to interrogate genetic risk for schizophrenia, we found that risk-associated enhancers in this disease are predominantly active in the brain, evolutionarily conserved, and AT-rich. The association between AT percentage and risk corresponds to an overrepresentation in risk-associated enhancers for the binding sites of transcription factors that recognize AT-rich cis-regulatory motifs. Several of the TFs identified in our model as being overrepresented in risk-associated enhancers, including MEF2C, are master regulators of neuronal development. The genes that encode several of these TFs are themselves located at genetic risk loci for schizophrenia. This list also includes brain-expressed TFs that have not previously been linked to schizophrenia. In summary, we developed a generalizable approach that integrates GWAS summary statistics with enhancer characteristics to identify risk factors in tissue-specific regulatory regions.

**AUTHOR SUMMARY:** Enhancers are regulatory regions that influence gene expression via the binding of transcription factors. Risk for many heritable diseases is enriched in regulatory regions, including enhancers. In this study, we introduce a novel method of testing for association between enhancer attributes and risk and use this method to determine the enhancer characteristics that are associated with risk for schizophrenia. We found that enhancers associated with schizophrenia risk are both evolutionarily conserved and in physical contact with mutation-intolerant genes, many of which have neurodevelopmental functions. Risk-associated enhancers are also AT-rich and contain binding sites for neurodevelopmental transcription factors.

## INTRODUCTION

### Regulatory regions in human disease

Non-coding genomic regions involved in gene regulation such as enhancers and promoters, as well as the transcriptional machinery that interacts with them, govern regulatory programs that underlie cell identity and specialization, patterning, and numerous functions necessary for the proper functioning of the body’s tissues and organs [1,2]. Genetic studies of many human traits indicate an enrichment of risk in these gene regulatory regions [3–8]. In genome-wide association studies (GWAS) of diseases such as cardiovascular, autoimmune, and neuropsychiatric disorders, more than 90 percent of SNPs in risk loci are non-coding variants [9]. Epigenomic studies over the past decade have mapped tissue- and cell type-specific gene regulatory elements in the non-coding genome, opening the door for large-scale exploration of their contribution to human disease. Such studies have demonstrated that disease-associated genetic variation is concentrated in regulatory regions in a tissue- and cell type-specific manner. For example, rheumatoid arthritis (RA) and Crohn’s disease risk are highly enriched in regions of accessible (active) chromatin from blood and immune cells, while type 2 diabetes risk is enriched in open chromatin from endocrine tissue [3]. Disease risk has also been connected to more specific regulatory elements, including enhancers and distal gene regulatory elements that activate and refine the cell type- and context-specific activity of many promoters [10].

These findings suggest that much of the genetic risk for complex traits acts through the disruption of regulatory regions unique to relevant tissue and cell types. However, there remain substantial gaps in our knowledge about the mechanisms by which variants in specific promoters and enhancers predispose to risk. This is in part because existing tools, while powerful, are not specifically designed to evaluate the features of specific regulatory regions that are associated with disease risk. Some of the most widely used approaches in this space include long distance interaction methods, fine-mapping tools, and approaches that leverage linkage disequilibrium. H-MAGMA performs gene-based association tests by aggregating the associations from all the SNPs in each gene’s distal regulatory regions predicted by Hi-C experiments to make inferences about genes, not regulatory elements [11]. Epigenomic fine-mapping tools such as PAINTOR and RiVIERA integrate non-coding annotations to predict specific, causal SNPs [12,13]. Stratified Linkage-Disequilibrium Score Regression (LDSC) is a highly effective tool for genome-wide inference of genomic features (e.g., open chromatin regions, evolutionarily conserved regions) enriched for disease risk, but is primarily used to assess binary annotations – rather than quantitative scores – and is underpowered for annotations representing less than ∼1% of the genome [3].

Here, we propose RWAS (for Regulome-Wide Association Study) to test associations of genetic risk with specific enhancers and enhancer properties. In the RWAS framework (Figure 1), we first collect enhancer annotations in a tissue relevant to the trait of interest, then identify risk-associated enhancers by aggregating the effects of all SNPs that overlap the enhancer’s position in the genome. Finally, we test associations of enhancer features with disease risk using a regression framework. RWAS is implemented as a novel application of MAGMA [9], which was originally developed for gene-based association studies and is widely used for that purpose. RWAS is fast, extensible, and requires only GWAS summary statistics and enhancer-level annotations and covariates. We apply RWAS to characterize enhancers and enhancer features that are associated with risk for schizophrenia (SCZ), a severe psychiatric disorder for which well-powered GWAS identified hundreds of risk loci enriched in brain-specific gene regulatory regions [9]. As part of this work, we also compiled a resource of high quality adult and fetal brain enhancer maps to identify risk-associated enhancer traits in the brain, suitable for association testing with RWAS. Our analyses reveal novel associations of SCZ risk with AT-rich enhancers in the developing brain as well as AT-rich transcription factor networks.

**Fig 1.**
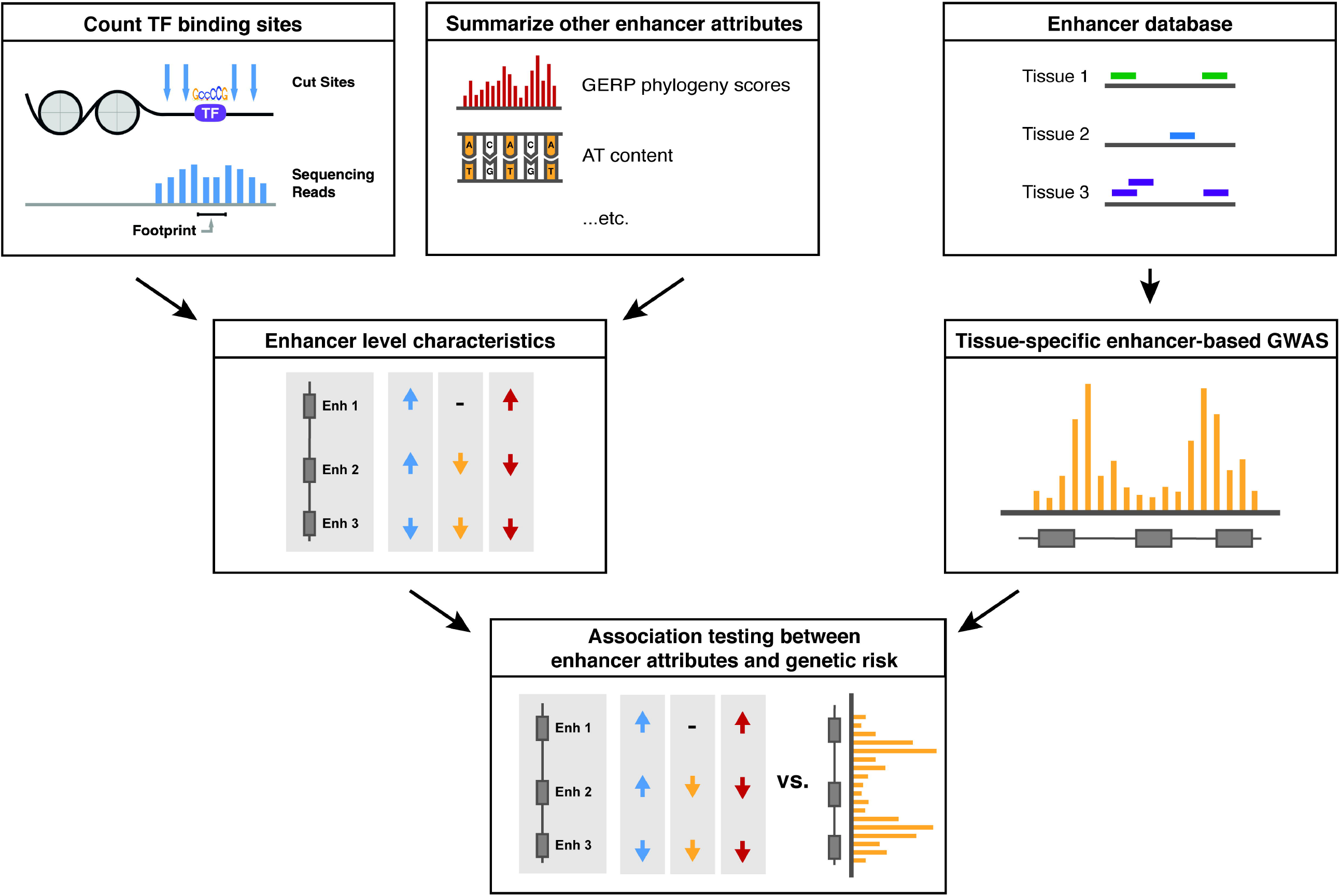
RWAS workflow overview. In brief, the RWAS workflow involves annotating SNPs to enhancers and other regulatory regions (rather than genes). Enhancer-level summary statistics are computed for input into association testing. Then, we use the MAGMA linear modeling framework to compute genetic associations between supplied enhancer-level covariates and these enhancer-based GWAS summary statistics. This approach relies on high-quality enhancer annotations for the tissue of interest that capture genetic risk for the disorder. To ensure these conditions were met, we first thoroughly characterized a set of brain-specific enhancers and demonstrated that these enhancers capture genetic risk for schizophrenia.

## RESULTS

### A database of enhancers and enhancer annotations in the human brain

The three elements required for an RWAS analysis are a database of tissue-specific gene regulatory elements, annotations describing the attributes of the enhancers to test for association testing, and GWAS summary statistics for a trait of interest. Here, we utilized chromHMM-derived enhancer predictions in 127 human tissues and cell types from the ROADMAP consortium as a regulatory element database [14]. These predictions are made using a hidden Markov model run on a series of epigenetic marks from each tissue, including chromatin accessibility and histone marks such as H3K27ac. A major advantage of the ROADMAP dataset is that the wealth of different tissues available makes cross-tissue comparison easier, enabling an unbiased view of enhancer activity across tissues and cell types. Analyses presented in this paper focus primarily on schizophrenia (SCZ), for which purpose we are primarily interested in annotations of enhancers in the brain. The dataset contains chromatin state annotations for 15 brain-related samples, including 7 samples from the adult brain, 3 from the prenatal brain at mid-gestation, 3 from embryonic stem cell (ESC)-derived neuronal progenitors or neurons, and 2 from neurosphere cultures. These annotations contain genomic coordinates of regions that were designated as enhancers by chromHMM’s hidden Markov model. In order to benchmark these enhancer annotations against other methods of enhancer annotation, we also studied predicted enhancer locations (regular- and high-confidence) from the psychENCODE consortium. These enhancers were derived from tissue from the prefrontal cortex, and were defined as areas enriched for acetylation at lysine 27 of the histone 3 tail (H3K27ac), which marks active regulatory regions, and depleted for tri-methylation at lysine 4 on the histone 3 tail (H3K4me3), which occurs primarily in promoters [15].

We validated these enhancer annotations by three different approaches. First, we compared enhancer locations in the 127 samples on the basis of summary statistics, including enhancer length, genomic coverage, enhancer number, and AT-richness (Figure S1). Brain enhancers were largely similar to other tissues in terms of size, length, and coverage (Figure S1A-F). Within the brain, adult brain samples had the highest coverage and number of predicted enhancers, while samples from fetal brain and models of neurodevelopment had lower coverage and number of enhancers (Figure S1A and B). Fetal brian and neurosphere samples had average enhancer lengths nearly 50 bp longer than those in adult brain samples (Figure S1C).

Second, we tested whether these enhancer annotations capture an element of tissue specificity. The Jaccard index was used to quantify pairwise similarity among the genomic locations of enhancers utilized in the 127 samples. As expected, enhancer utilization clustered samples by organ, as well as by developmental age (Figure S3). In the brain, we found three groups of samples distinguished by their enhancer utilization, corresponding to adult brain, fetal brain and cultured neurospheres, and cultured neural progenitors (Figure 2).

**Fig 2.**
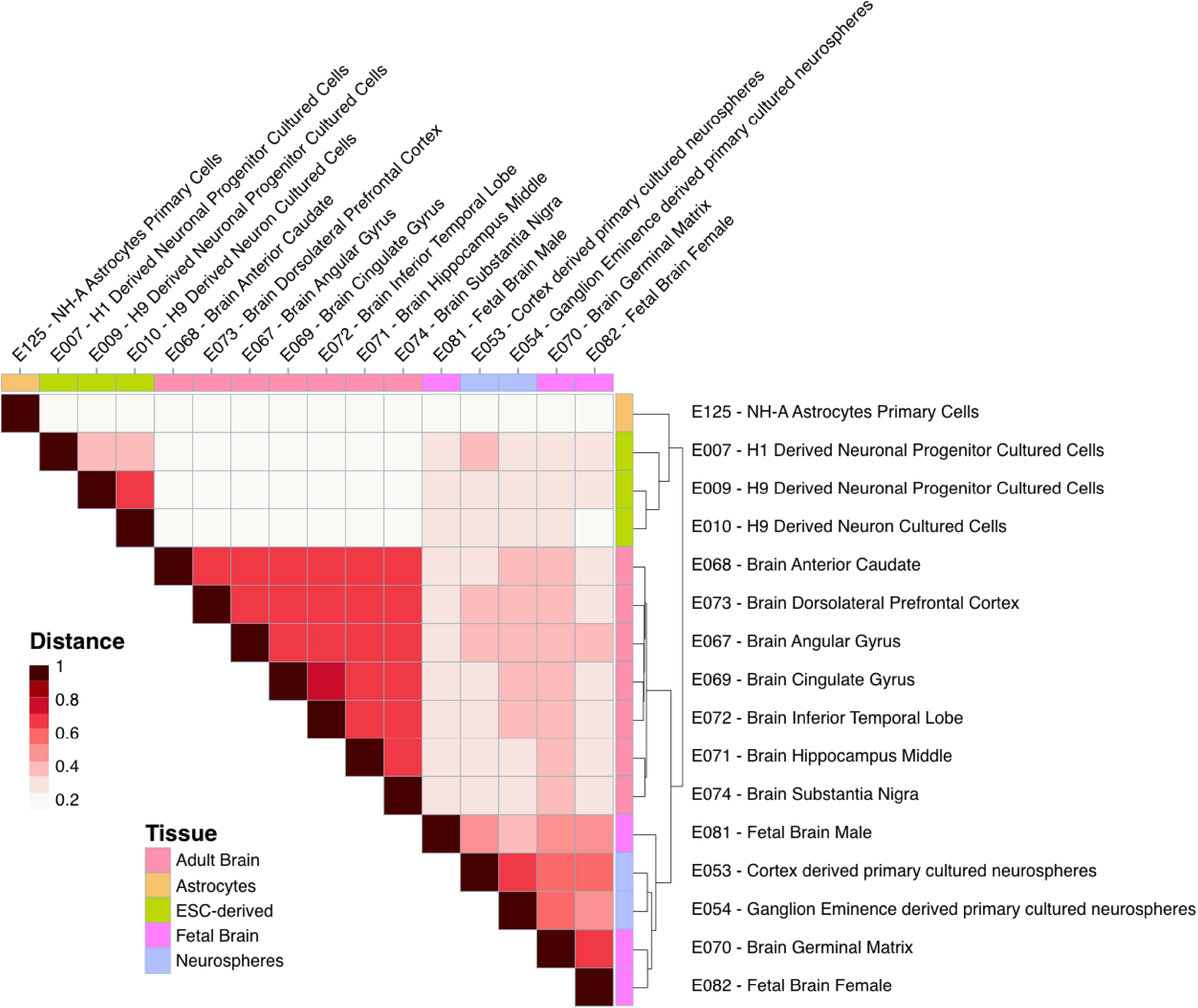
Genome-level Jaccard similarity matrix demonstrates age- and experimental model-specific enhancer patterning. Fetal brain and neurosphere samples cluster together when the tree is cut at the second level, while adult brain samples, ESC-derived clusters, and astrocytes form separate groups. Color denotes Jaccard distance statistic. Groupings determined using hierarchical clustering.

Third, we tested that our enhancer annotations confirm known associations, focusing on SCZ [5]. Previous studies have shown that enhancers and other gene regulatory regions active in the human brain are enriched for heritability in SCZ [3–5]. As expected, stratified LD Score Regression using summary statistics from schizophrenia GWAS [5] confirmed that brain enhancers from our analysis were highly enriched for SCZ risk (Figure 3A). The adult brain-specific enhancer annotation most significantly enriched for SCZ risk was the inferior temporal gyrus (sample E072, p= 4.6E-14), and the fetal brain-specific enhancer annotation most significantly enriched for risk was female fetal brain (sample E082, p= 1.28E-9). These enrichments were comparable in significance to the enrichment of SCZ risk in two sets of adult prefrontal cortex enhancers from the PsychENCODE consortium (Figure 3B) [15]. In summary, our validation tests indicate that ROADMAP ChromHMM models provide robust annotations of enhancers in the fetal and adult brain that capture a tissue-specific element of genetic risk for SCZ. These analyses define a total of 388,011 non-overlapping enhancer regions and are available at http://data.nemoarchive.org/other/grant/sament/sament/RWAS.

**Fig 3.**
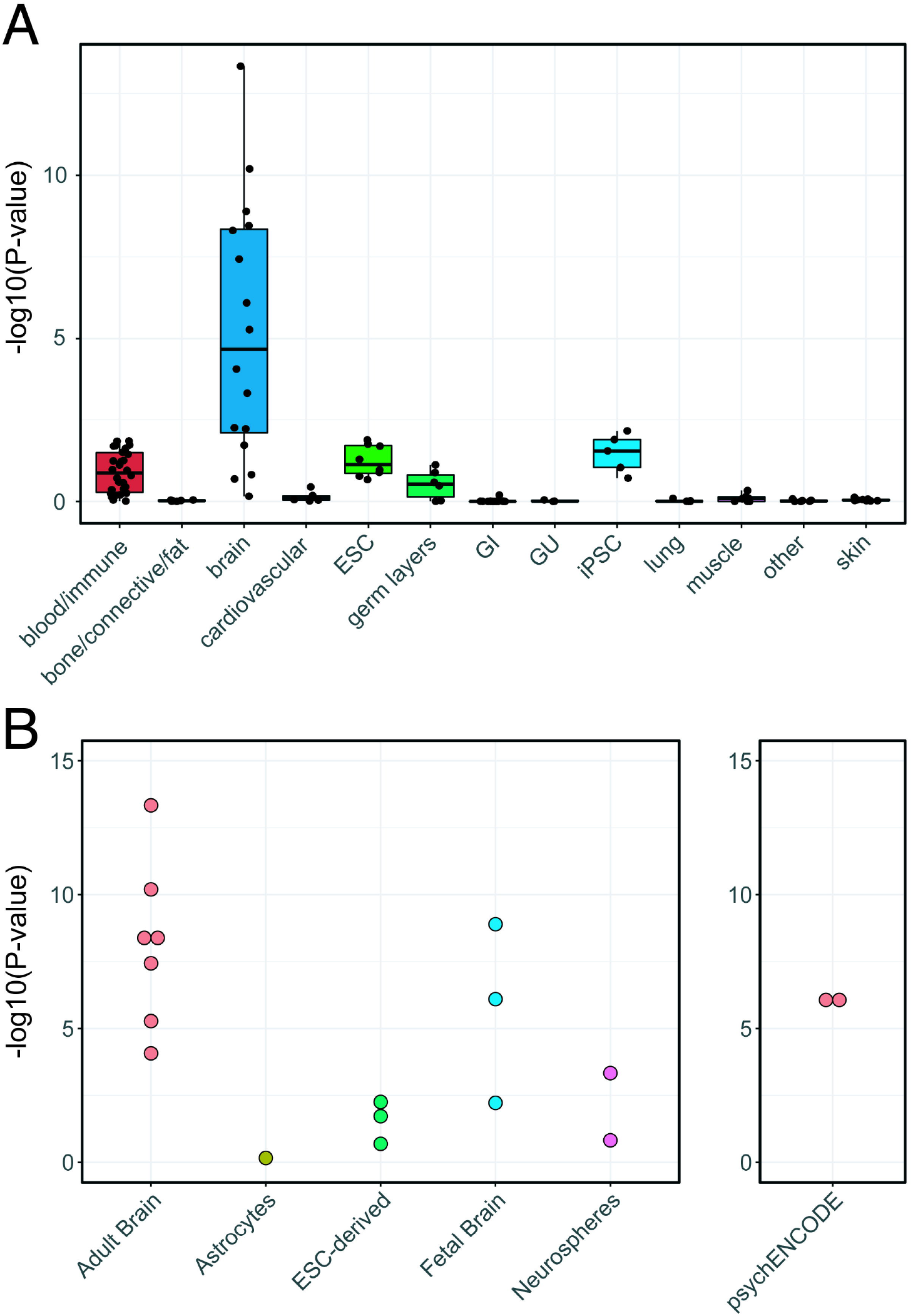
Genetic risk for schizophrenia is enriched in chromHMM-derived brain enhancers. A) Partitioned heritability of enhancer annotations by tissue in schizophrenia. Brain enhancers are enriched for heritability in schizophrenia compared to other tissues. B) Partitioned heritability of individual brain samples. Adult brain enhancers had the most significant enrichment, followed by fetal brain enhancers.

### RWAS reveals enhancers and enhancer characteristics associated with risk for schizophrenia

We hypothesized that SCZ risk is associated with SNPs that impact specific enhancers that are active in the brain. To identify these enhancers, we performed an “enhancer-based” GWAS analysis of the PGC2 SCZ GWAS, testing for significance for the aggregated SNPs within each enhancer using the SNP-wise regression model implemented in MAGMA (Figure 4A). This analysis revealed a total of 2,784 risk-associated enhancers at a genome-wide significance threshold p < 1.3E-7, which corresponds to alpha < 0.05 after Bonferroni correction for 388,011 non-overlapping brain-activated enhancer regions in our database (Figure 4B). 2,001 of these risk-associated brain enhancers are located within 63 of the 108 risk loci identified in the original (SNP-based) analysis of these data, while the remaining enhancers are at loci that did not reach genome-wide significance in the primary analysis. Examination of specific risk loci indicated that risk-associated enhancers capture the genetic risk signal at many of the SNP-based risk loci in a tissue-specific manner (Figure 4B). These findings demonstrate that tissue-specific enhancer based GWAS can capture risk signal from schizophrenia GWAS summary statistics.

**Fig 4.**
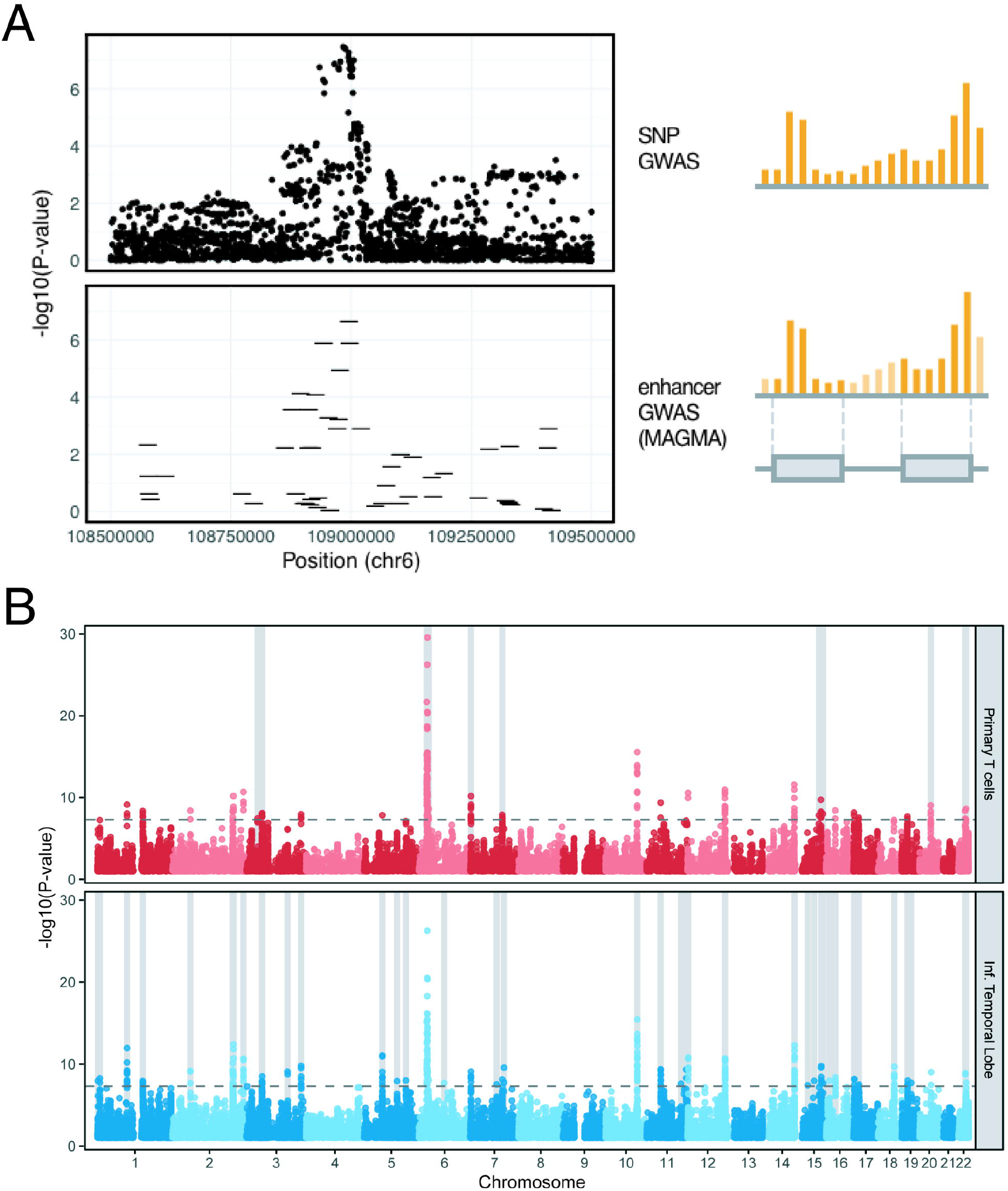
Enhancer-based RWAS demonstrates tissue specificity of genetic risk capture. A) Enhancer-based GWAS allows the aggregation of non-coding SNPs into nearby enhancer regions and captures risk loci as enhancer-level risk associations. A short region of chromosome 6 containing a risk locus is shown as an example. Individual points on the top panel denote SNPs, while the lines on the bottom panel represent enhancer regions. B) Brain enhancers capture risk loci missed by enhancers from other tissues. Enhancers from primary T cells captured fewer genome-wide significant SNPs when compared to enhancers from the inferior temporal lobe of the adult brain. The light gray shaded areas denote loci where the enhancer annotation has more genome-wide significant SNPs when compared to the other annotation.

For validation, we integrated the SCZ-associated enhancers from our analysis with results from massively parallel reporter assays (MPRA) of schizophrenia risk alleles [16] and with expression quantitative trait loci (eQTLs) in the prefrontal cortex [17]. This analysis provided independent evidence for several of the top risk-associated enhancers in our analysis. A fetal brain-specific enhancer at chr1:243555100-243556100 (p= 7.68E-11), located in an intron of *SDCCAG8*, contains a SNP (rs77149735) associated with differential enhancer activation by MPRA. A second fetal brain-specific, risk-associated enhancer from our analysis, located at chr22:42657000-42658000 (p=2.4E-9) contained the SNP rs134873, which was significantly associated with differential enhancer activity in MPRA assays and an eQTL for the genes FAM109B, NAGA, LINC00634, and WBP2NL. These results demonstrate both that our enhancers capture variants that have strong evidence of regulatory impact and that our enhancer-based association tests can be integrated with other datasets to identify regulatory regions for further follow up.

The RWAS framework enables enhancer-property analyses to test for association between disease risk and a variety of enhancer features. Applying this approach to enhancers associated with risk for SCZ, we first tested the hypothesis that risk-associated enhancers regulate gene sets that have previously been implicated in neuropsychiatric studies. We used Hi-C data from the developing brain [18] to predict the targets of risk-associated enhancers. Across all the adult and fetal brain enhancer annotations, a total of 720 genes were in contact with at least one statistically significant risk-associated enhancer. These associations were quite reproducible: 648 of these genes were identified in more than one brain tissue enhancer annotation and 248 were found in all ten. Using these enhancer-gene maps, we tested for enrichments in 64 gene sets that have previously been implicated in SCZ risk (Figure 5A). Enhancer targets were strongly enriched for genes that are intolerant of loss-of-function mutations (p=2.37E-3, pLI; p=2.16E-4, LOEUF [see methods for definition]). Risk-associated enhancers also disproportionally contact genes that are bound by the neuron-specific RNA-binding proteins Fragile X mental retardation protein (FMRP) (p = 3.07E-5) and RBFOX1/3 (p = 3.75E-4), as well as targets of the autism-associated chromatin remodeling gene Chromodomain-helicase-DNA-binding protein 8 (CHD8) (p = 3.99E-4). Rare mutations in FMRP and CHD8 cause neurodevelopmental disorders with autistic features [19–26] and regulate neurodevelopmental gene networks that have previously been linked to SCZ in genetic and proteomic studies [27,28]. These findings extend previous gene-based analyses [29,30].

**Fig 5.**
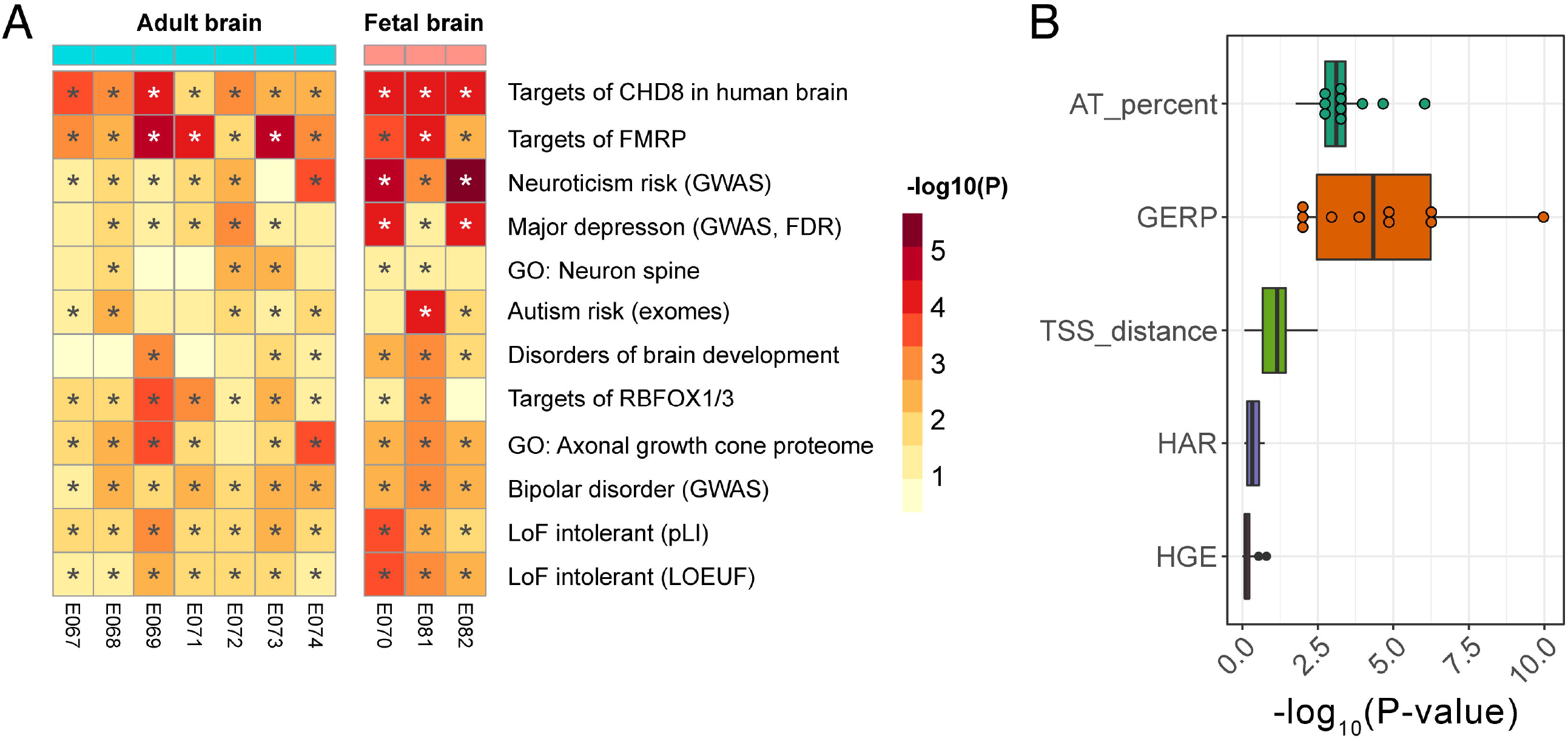
A) Risk-associated enhancers are in physical contact with mutation intolerant genes and targets of RNA-binding proteins with critical roles in development. RWAS reveals target genes and enhancer characteristics associated with schizophrenia risk. GWAS = genes at GWAS risk loci. Full description of gene lists available in Methods. B) Association testing between schizophrenia risk and enhancer attributes. High GERP scores and high AT percentages at the enhancer level were independently associated with schizophrenia risk. HGE/HAR status and distance to the nearest gene TSS were not associated with risk.

Next, we tested the hypothesis that risk-associated enhancers differ in their evolutionary history from other enhancers in the brain. Enhancers with deep evolutionary conservation may have particularly important functions in the brain. It has also been postulated that risk for SCZ may involve evolutionarily novel enhancers, some of which regulate human-specific aspects of brain development [31,32]. Evolutionary conservation within enhancer regions (defined by GERP phylogeny scores) was positively associated with risk (Figure 5B), with fetal brain enhancers having the most significant associations (male fetal brain, 1.1E-10; germinal cortex at 20wk gestation, 5.2E-7; female fetal brain, 5.9E-7; all 10 adult and fetal brain enhancer annotations significant at FDR < 0.05). By contrast, enhancers in two categories of evolutionarily novel enhancers – human accelerated regions (HARs) and human-gained enhancers (HGEs) – were not significantly associated with schizophrenia risk (Figure 5B), in agreement with previous results [21]. This finding is unlikely to be due to low power, since 12,501 brain enhancers were found within 5kb of an HGE and 7,984 were located within a HAR. Therefore, schizophrenia risk-associated enhancers are older in evolutionary time and are not generally under positive selection.

Since many enhancers regulate proximal promoter regions, we hypothesized that enhancers closer to a transcription start site would be more strongly associated with disease risk. However, we found that distance to the nearest gene was not associated with risk (Figure 5B). This is in line with the discovery of significant long-range interactions between schizophrenia risk SNPs and genes with neuronal functions [18].

Unexpectedly, one of the enhancer features most strongly associated with SCZ risk was the percent of adenine-thymidine base pairs (AT richness), which was positively associated with risk across all brain tissues surveyed (Figure 5B). The most significant association in adult brain was in prefrontal cortex enhancers (E073, p = 9.1E-7), while the strongest association in the fetal brain was in the fetal germinal matrix (E070, 20 weeks gestational age; p = 5.68E-6). Overall, brain enhancers do not have substantially higher AT richness than enhancers in other tissues (Figure S2C). In addition, we did not find a strong association between AT richness and SCZ risk within enhancers from other tissues (Figure S4). Therefore, these results suggest that SCZ risk is enriched specifically at AT-rich enhancers in the adult and developing brain.

### SCZ-associated enhancers are enriched for binding sites for neurodevelopmental transcription factors recognizing AT-rich sequence motifs

We hypothesized that the association of AT risk enhancers with SCZ corresponds with occupancy by transcription factors that recognize AT-rich sequence motifs. To test this, we performed an RWAS testing for association between SCZ risk and binding sites for individual TFs. We used tissue-specific TF binding site predictions for 503 TFs, derived from integration of DNase-seq footprinting analysis in the human brain with JASPAR2016 vertebrate sequence motifs [33]. There was a strong association between the AT-richness of a given TF binding site motif and the effect size in our model (p= 2.0E-15 in female fetal brain). These associations were also borne out in a meta-analysis performed by combining all 10 adult and fetal brain enhancer RWAS (p < 2E-16). We also found that TF motifs with positive association with SCZ risk in our RWAS had a higher AT percentages than motifs with a negative association; in other words, TF motifs that were overrepresented in risk-associated enhancers had higher AT percentages than TF motifs that were depleted in risk-associated enhancers (Figure 6A).

**Fig 6.**
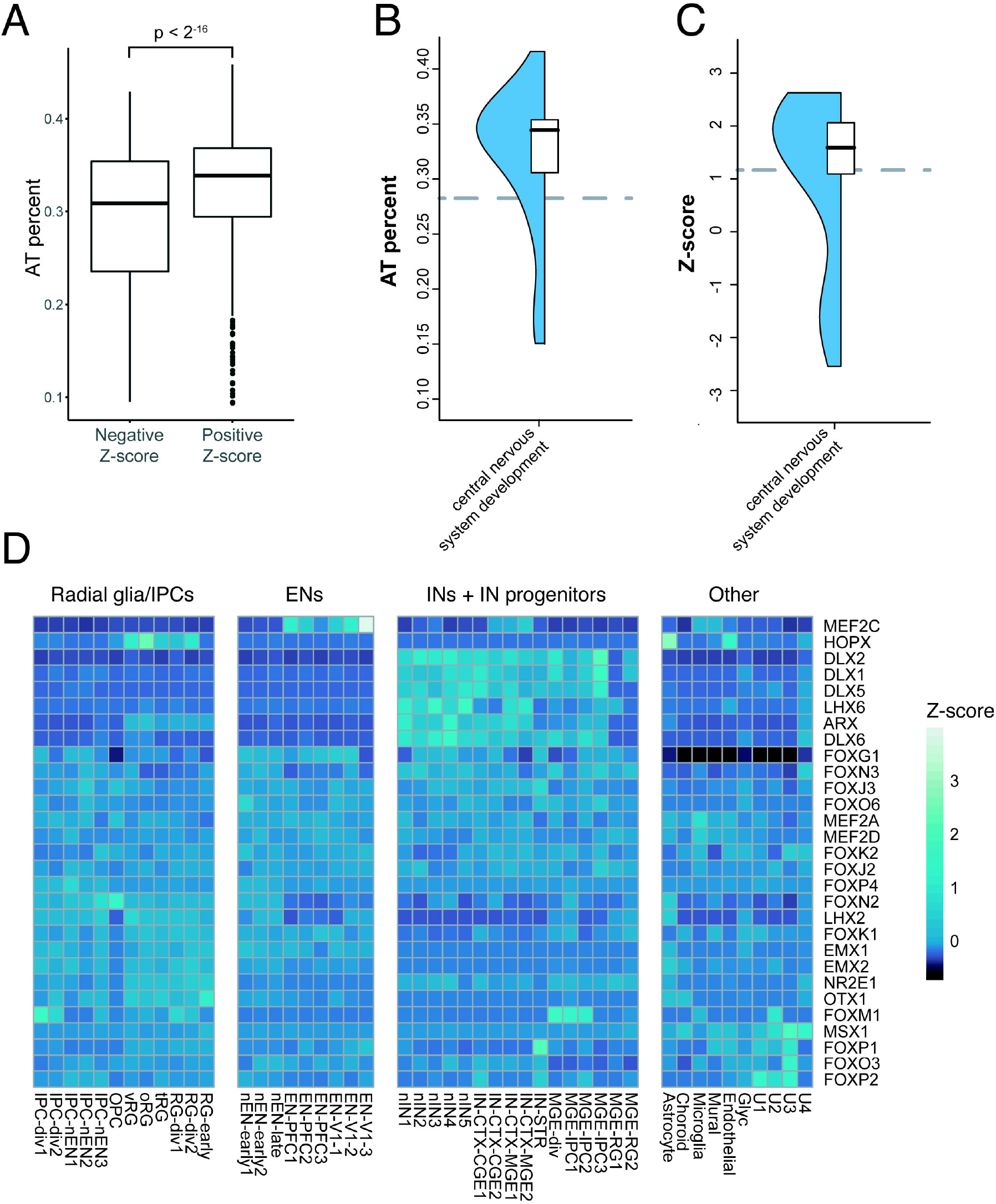
TFs with AT rich motifs are overrepresented in risk-associated enhancers and have neurodevelopmental function. A) There is a strong association between the AT-richness of a given TF binding site motif and the direction of the effect size in the RWAS model. AT rich motifs tend to have a positive association with risk. B) Higher median motif AT percentage of a given transcription factor is positively associated with the TF playing a role in cell morphogenesis during neuron differentiation. C) TFs with higher median Z-score in the RWAS analysis are more likely to play a role in cell morphogenesis during neuron differentiation. Grey dashed line is the median value of the background set of all TFs in our dataset. D) TFs that recognize positively associated motifs in the schizophrenia RWAS analysis are involved in neural development. Each cell is colored by scaled expression averaged across the specified cell type. The TFs shown are expressed in > 250 cells in the developing brain and have at least one motif with a meta-analyzed z-score of > 3 in the RWAS model. EN = excitatory neuron, IN = inhibitory neuron.

While the nucleotide composition of promoters and of larger chromosomal segments (isochores, >300 kb on average) has been extensively described, the functional differences between AT-rich vs. GC-rich enhancers are not well understood. Strikingly, many of the most positively associated sequence motifs, all of which are AT rich, are recognized by neurodevelopmental TFs, including members of the MEF2 family, the EMX family, and the DLX family (Table 1). Based on this result, we asked whether AT-richness might be a general feature of neurodevelopmental TFs. Indeed, TFs annotated to the Gene Ontology term “cell morphogenesis involved in neuron differentiation” and related GO terms had higher motif AT percentages than other TFs (Wilcoxon p = 5.0E-5, Figure 6B), and motifs recognized by these neurodevelopmental TFs were positively associated with SCZ risk in our model (p = 1.64E-3, Figure 6C).

**Table 1.**
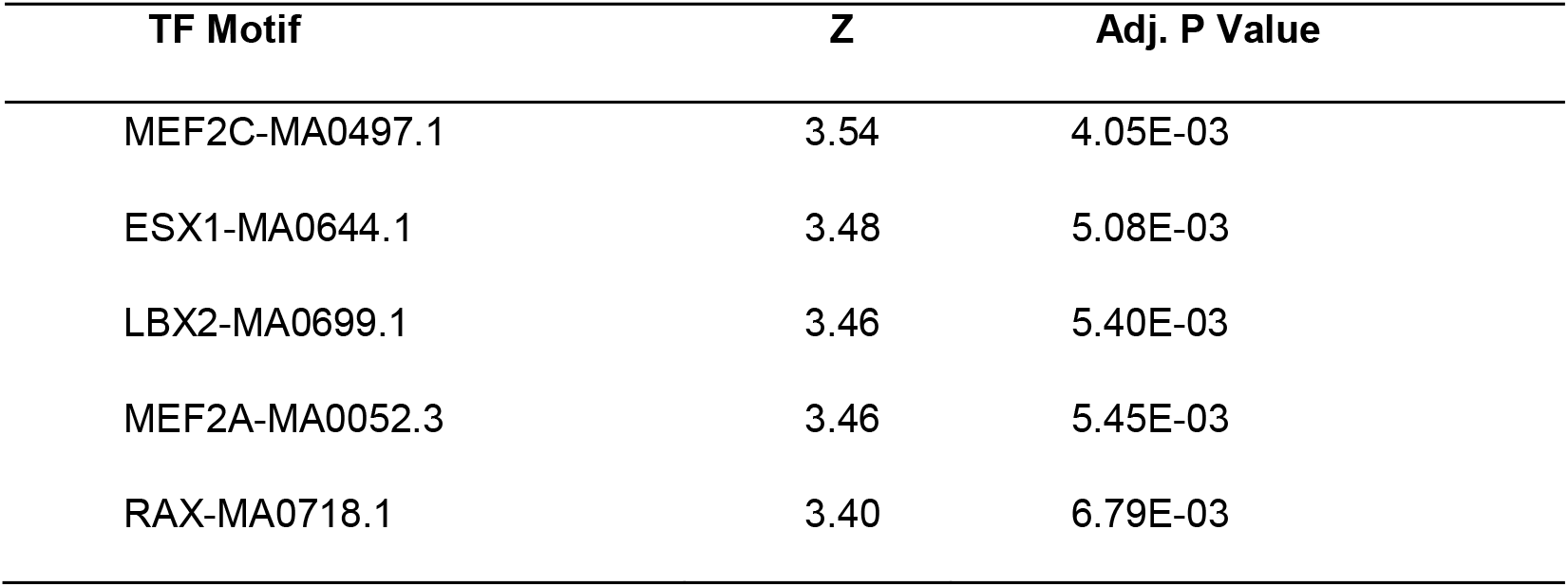
Top 5 strongest positive associations between TF motif networks and schizophrenia risk in brain enhancers. While these motifs are recognized by a variety of TFs, a few families that utilize these motifs are of particular interest in the brain: for example, the MEF2C motif is recognized by all members of the MEF family and the LBX2 motif is recognized by the DLX, EMX, and ARX families.

| TF Motif | Z | Adj. P Value |
| --- | --- | --- |
| MEF2C-MA0497.1 | 3.54 | 4.05E-03 |
| ESX1-MA0644.1 | 3.48 | 5.08E-03 |
| LBX2-MA0699.1 | 3.46 | 5.40E-03 |
| MEF2A-MA0052.3 | 3.46 | 5.45E-03 |
| RAX-MA0718.1 | 3.40 | 6.79E-03 |

We further explored the developmental expression patterns of TFs that recognize SCZ-associated sequence motifs using single-cell RNA sequencing data from prenatal human cortex [34]. We found that many of the TFs that are highly associated with risk in our model are expressed in neuronal lineages, including members of the MEF2 family, the EMX family, the RAX family, and the DLX family (Figure 6D). Taken together, these results suggest a previously undescribed association between SCZ risk and AT-rich binding sites for neurodevelopmental TFs in enhancers of the fetal and adult brain.

## DISCUSSION

Here, we developed tools and resources for Regulome-Wide Association Studies (RWAS), a flexible framework for post-GWAS analyses of trait-associated enhancers and enhancer properties. Using these tools, we characterized enhancers associated with risk for schizophrenia.

Our analysis revealed a novel association of SCZ with AT-rich enhancers that are active in the human brain, many of which contain AT-rich sequence motifs recognized by neurodevelopmental TFs. Functional differences between AT-rich vs. GC-rich enhancers are not well understood. One previous study using Cap Analysis of Gene Expression showed that many enhancers actively transcribed in neurons are AT-rich and noted differences in TF occupancy in GC-rich vs AT-rich enhancers [35]. Our analysis generalizes this observation to a broader set of enhancers defined by independent epigenomic techniques. Functional differences of AT-rich vs. GC-rich promoters are better characterized, with AT-rich promoters containing distinct core promoter elements and serving different functions. For example, Lecellier et al. demonstrated that AT-rich promoter regions were disproportionately found near genes involved in the immune response [36]. Large genomic regions of relatively consistent nucleotide composition in the genome, known as isochores, have also been described to contain genes with shared functions; for example, GC-rich isochores tend to contain housekeeping genes, while AT-rich isochores tend to contain more tissue-specific genes [37]. To our knowledge, we are the first to report that neurodevelopmental TFs predominantly recognize AT-rich sequence motifs.

The specific neurodevelopmental TFs whose putative binding sites were enriched at SCZ-associated enhancers represent promising leads toward mapping the causal gene regulatory perturbations underlying SCZ. The most significant positive association in our TF RWAS was MEF2C-MA0497.1. This association is consistent with previous reports that the MEF2C motif is enriched at SCZ risk loci, and MEF2C target genes in the brain are enriched both for SCZ risk genes [38,39] and for genes differentially expressed in postmortem brain tissue from SCZ cases vs. controls. *MEF2C* itself is a positional candidate at an SCZ risk locus [5]. MEF2C is highly expressed in developing cortical excitatory neurons and is essential both for cortical neurogenesis and the modulation of cortical neuronal activity. Haploinsufficiency of MEF2C is known to cause a syndrome characterized by intellectual disability and neurological abnormalities [40]. Another network of interest is LBX2-MA0699.1, a motif recognized by multiple homeobox TFs. Of particular interest are EMX1 and EMX2, which are highly expressed in the developing dorsal telencephalon in the lineage leading to excitatory neurons and have well-established roles in cortical thickness and arealization [41–44]. Similarly to MEF2C, the area containing the gene for EMX1 is itself a candidate schizophrenia risk locus [5]. Mutations in EMX2 have been noted in patients with severe schizencephaly [27]. The LBX2-MA0699.1 motif is also recognized by members of the DLX and ARX families. Unlike the EMX factors that are involved in excitatory neuron development, these TFs are critical for inhibitory neuron development and migration [28–30]. Mutations in ARX have been linked to cases of X-linked lissencephaly with abnormal genitalia in humans [30]. A limitation of our analysis is that motif-based predictions cannot resolve the specific members of this TF family that occupy the SCZ-associated enhancers, but the family as a whole merits increased attention in SCZ.

RWAS is readily applicable to additional traits of interest, as it is implemented with a widely used software tool (MAGMA) and requires only GWAS summary statistics and enhancer level annotations. We have made our instructions for running RWAS available at www.github.com/casalex/RWAS. We have made available the enhancer annotations for all 127 ROADMAP samples, and similar enhancer models suitable for RWAS are now available from >800 samples from the ENCODE consortium, spanning all of the major human organs and tissues [45].

## METHODS

### Enhancer download and processing

Predicted enhancer regions were derived from 25-state ChromHMM [14] chromatin state models downloaded from the ROADMAP consortium website (https://egg2.wustl.edu/roadmap/web_portal/imputed.html). We defined enhancers by pooling nine states from these models: 1) transcribed 5’ preferential and enhancer, 2) transcribed 3’ preferential and enhancer, 3) transcribed and weak enhancer, 4) active enhancer 1, 5) active enhancer 2, 6) active enhancer flank, 7) weak enhancer 1, 8) weak enhancer 2, and 9) primary H3K27ac possible enhancer. Enhancer annotations from the psychENCODE consortium were downloaded from http://resource.psychencode.org/. The brain enhancers used in the schizophrenia RWAS analyses were E067 (Brain Angular Gyrus), E068 (Brain Anterior Caudate), E069 (Brain Cingulate Gyrus), E070 (Brain Germinal Matrix), E071 (Brain Hippocampus Middle), E072 (Brain Inferior Temporal Lobe), E073 (Brain_Dorsolateral_Prefrontal_Cortex), E074 (Brain Substantia Nigra), E081 (Fetal Brain Male), and E082 (Fetal Brain Female).

Enhancer annotations used for partitioned heritability and RWAS analyses were pre-processed in a uniform pipeline. Enhancer boundaries are often poorly defined, and MAGMA and similar tools suffer from length bias wherein long regions with many SNPs have anti-conservative p-values (Figure S5). To overcome these issues, our analyses were conducted using 1 kb enhancer centroids. Enhancer regions were merged with any directly adjacent annotations, and the center of each merged region was determined. The boundaries were then extended by 500 bp upstream and downstream of this center, resulting in a 1kb region centered on the middle of the enhancer region. Any enhancers falling within the MHC region or ENCODE blacklist regions [46] were removed.

### Jaccard similarity

In order to compare enhancer similarity across all 127 samples we computed pairwise genome-wide Jaccard distance using the BEDtools software suite [47]. Groupings were determined using hierarchical clustering.

### GWAS summary statistics

We retrieved GWAS summary statistics for schizophrenia [5] from the Psychiatric Genomics Consortium data portal (https://www.med.unc.edu/pgc).

### Partitioned heritability

Stratified LD score regression (LDSC version 1.0.1) was applied to GWAS summary statistics to evaluate the enrichment of trait heritability across the 127 enhancer sets [33]. These associations were adjusted for 52 annotations from version 1.2 of the LDSC baseline model (including genic regions, enhancer regions and conserved regions).

### RWAS

RWAS was performed using the linear model implemented in MAGMA’s covariate mode. This was accomplished by using the enhancer sets in place of genes. The processed enhancers were supplied as a genomic location file format as described in the MAGMA manual, and enhancer-level attributes were supplied as continuous covariates. GERP hg19 phylogeny scores were downloaded from http://hgdownload.cse.ucsc.edu/goldenpath/hg19/phastCons100way/ and averaged across each enhancer region to yield a conservation score for each enhancer for association testing. TSS for each gene were taken from a supplied MAGMA gene file (https://ctg.cncr.nl/software/magma). Distance to the nearest TSS for each enhancer was determined using the BEDTools “closest” command, and this distance was supplied to MAGMA as a covariate for association testing. HAR regions were downloaded from Supplemental Table 1 of Doan et al. (2016) [48]. These regions were expanded by 2,500 bp upstream and downstream before being intersected with the enhancer regions, yielding a binary measure for each enhancer indicating if an enhancer overlapped an HAR or not. This was input as a covariate in the MAGMA analysis. Similarly, HGEs were defined as differentially enriched CREs between human and rhesus macaque from Vermunt et al. and overlaps were tested for association [49].

Chromosomal contact testing was performed using the set analysis in MAGMA. We used HiC from the cortical plate of the developing human brain [18] to assign genes to enhancers that they physically contact. Enhancers that contact genes with a given ontology term were assigned to the enhancer set for that term, and the resultant enhancer sets were tested for association with risk using MAGMA’s gene set mode. The gene sets are available in Supplementary Table 1 and are derived from the following datasets: genes intolerant of loss-of-function variants from gnomAD (pLI >= 0.9 or LOEUF deciles 1 or 2) [50]; risk genes from studies of rare variants in four disorders, including severe developmental disorder risk genes from the Deciphering Developmental Disorders consortium’s DDG2P database (Disorders of Brain Development) [51], autism spectrum disorder risk genes from the Autism Sequencing Consortium (Autism risk [exomes]) [52], bipolar disorder risk genes from the BipEx Consortium [53]; genes identified from large-scale GWAS, identified by gene-based analyses with MAGMA [9] (p < 2.77e-6 unless noted as FDR, in which case adj. p < 0.05) for bipolar disorder [54], major depression [55], and neuroticism [56], differentially expressed genes in the prefrontal cortex of individuals with schizophrenia, bipolar disorder, and autism from the PsychENCODE consortium [57] (http://resource.psychencode.org/Datasets/Derived/DEXgenes_CoExp/DER-13_Disorder_DEX_Genes.csv); genes associated with schizophrenia from SCHEMA [58] (25 genes with SCHEMA P < 0.05; odds ratio = 1.6, P = 0.03); target gene networks of the neuropsychiatric risk genes FMRP, RBFOX1/3, RBFOX2, CHD8, CELF4, and microRNA-137 derived from functional genomics experiments, annotated by Genovese et al. [59]; synaptic genes from SynaptomeDB (“GO: Axonal growth cone proteome, GO: Neuron spine”) [60]. In-text p-values were derived by taking the minimum p-value across the 10 brain enhancer annotations and adjusting for the number of annotations.

### TF binding site RWAS

Brain-specific DNAse-seq footprints annotated with matching TF motifs were obtained from our previously described footprint atlas [33]. The HINT atlas was used due to its superior performance in TF binding site prediction. A HINT score cutoff of 55 was used to filter out low-quality footprints. We limited our analysis to the 503 JASPAR vertebrate core motifs that had mappings to human TFs. Footprints that fell within the boundaries of a given enhancer were annotated to that element, yielding a covariate file containing counts of each motif for each enhancer. A total binding site control was used to control for total binding site number. MAGMA was run in the covariate mode as described above. Meta-analysis of adult and fetal brain enhancer RWA analyses was performed by taking the highest absolute value Z-score from the individual enhancer RWA results for each motif. The resultant p-values were adjusted for the number of results meta-analyzed (10).

### Motif to TF mapping

The footprint-motif pairs were mapped to TFs using a key described in our previous work [33]. These mappings were restricted to JASPAR motifs, so only these motifs were included in downstream analyses.

### GO term analysis

We used the Wilcoxon rank-sum test as implemented in the R package GOfuncR to test for association between TF function and scores/attributes from our models.

### Single-cell RNA TF expression

The single-cell RNA-seq dataset from the prenatal human cortex was downloaded from the UCSC cell browser (http://cells.ucsc.edu/cortex-dev/exprMatrix.tsv.gz) [34]. The Z-scores were generated using R’s scale() function, grouped by cell type, then averaged.

## Data Availability

We have made ready-to-use MAGMA-formatted enhancer location files for all 127 annotations used in this paper and instructions on creating covariate files publicly available at www.github.com/casalex/RWAS.

## Acknowledgements

We would like to thank Peter Zandi and Pippa Thompson for furnishing us with the gene lists used in Figure 5A. We would also like to acknowledge our funding sources: NIMH F30 MH120910 (PI: AMC) and NIMH R24MH114815 (PI: Hertzano). We would also like to thank the University of Maryland Medical Scientist Training Program for their ongoing assistance and mentorship.

## Supporting information captions

Fig S1. Enhancer annotation summary statistics. A) Genomic coverage by tissue category. B) Adult brain and astrocyte enhancer annotations had the highest genomic coverage compared to fetal, neurosphere, and ESC-derived enhancer annotations. C) Enhancer number by tissue category. D) Adult brain and astrocyte enhancer annotations had the highest enhancer number compared to fetal, neurosphere, and ESC-derived enhancer annotations. E) Enhancer length by tissue category. F) Fetal brain and neurosphere enhancer annotations had the highest mean enhancer length compared to adult brain, astrocyte, and ESC-derived enhancer annotations. G) Enhancer number and percent genomic coverage are tightly associated (p=8.1E-59) . H) Enhancer length and genomic coverage are not associated (p=0.64) . I) Enhancer length and enhancer number are negatively correlated (p=9.1E-6).

Fig S2. Enhancer annotations vary by length and nucleotide composition. A) Fetal brain and neurosphere enhancer annotations are underrepresented in the lowest length bins compared to other brain samples. B) Fetal brain and neurosphere enhancer annotations have more super-long enhancers than other brain enhancer annotations. C) AT percentage of enhancer annotations by tissue. D) Adult brain enhancer annotations have slightly higher AT richness compared to fetal brain and neurosphere enhancer annotations. E-F) Super-long enhancers tend to be more AT rich than regular enhancers.

Fig S3. Jaccard distance between all 127 chromHMM enhancer annotations. Annotations are arranged by hierarchical clustering. Differential enhancer utilization between tissues clusters samples by organ. Brain enhancers largely cluster together, with the exception of ESC-derived cells and astrocytes.

Fig S4. Association between AT-richness and schizophrenia risk across all chrommHMM enhancers aggregated by tissue. Brain enhancers had the strongest association, with germinal matrix (E070) having the most associated individual annotation.

Fig S5. Non-truncated enhancers suffer from length bias in MAGMA gene-set analyses. A. Z-scores are higher in longer enhancers compared to shorter enhancers in the PGC2 schizophrenia GWAS. B. Z-scores show similar inflation in long enhancers in 75 unrelated UK Biobank traits.

## Notes

### Competing Interest Statement

The authors have declared no competing interest.

## REFERENCES

1. Davidson EH. Gene Regulatory Networks for Development. The Regulatory Genome. Elsevier; 2006. pp. 125–185. doi:10.1016/b978-012088563-3.50022-5

2. Dunham I, Kundaje A, Aldred SF, Collins PJ, Davis CA, Doyle F, et al. An integrated encyclopedia of DNA elements in the human genome. Nature. 2012;489: 57–74. doi:10.1038/nature11247

3. Finucane HK, Bulik-Sullivan B, Gusev A, Trynka G, Reshef Y, Loh P-R, et al. Partitioning heritability by functional annotation using genome-wide association summary statistics. Nature genetics. 2015;47: 1228–35. doi:10.1038/ng.3404

4. Finucane HK, Reshef YA, Anttila V, Slowikowski K, Gusev A, Byrnes A, et al. Heritability enrichment of specifically expressed genes identifies disease-relevant tissues and cell types. Nature Genetics. 2018;50: 621–629. doi:10.1038/s41588-018-0081-4

5. Ripke S, Neale BM, Corvin A, Walters JT, Farh K-H, Holmans PA, et al. Biological Insights From 108 Schizophrenia-Associated Genetic Loci. Nature. 2014;511: 421–427. doi:10.1038/nature13595

6. Maurano MT, Humbert R, Rynes E, Thurman RE, Haugen E, Wang H, et al. Systematic localization of common disease-associated variation in regulatory DNA. Science. 2012;337: 1190–1195. doi:10.1126/science.1222794

7. The GTEx Consortium, Welter D, MacArthur J, Morales J, Burdett T, Hall P, et al. The Genotype-Tissue Expression (GTEx) pilot analysis: multitissue gene regulation in humans. Science. 2015;348: 648–60. doi:10.1126/science.1262110

8. Freudenberg J, Gregersen P, Li W. Enrichment of Genetic Variants for Rheumatoid Arthritis within T-Cell and NK-Cell Enhancer Regions. Molecular medicine (Cambridge, Mass). 2015;21: 180–4. doi:10.2119/molmed.2014.00252

9. de Leeuw CA, Mooij JM, Heskes T, Posthuma D. MAGMA: generalized gene-set analysis of GWAS data. PLoS computational biology. 2015;11: e1004219. doi:10.1371/journal.pcbi.1004219

10. Claussnitzer M, Dankel SN, Kim K-H, Quon G, Meuleman W, Haugen C, et al. FTO Obesity Variant Circuitry and Adipocyte Browning in Humans. The New England journal of medicine. 2015;373: 895–907. doi:10.1056/NEJMoa1502214

11. Sey NYA, Hu B, Mah W, Fauni H, McAfee JC, Rajarajan P, et al. A computational tool (H-MAGMA) for improved prediction of brain-disorder risk genes by incorporating brain chromatin interaction profiles. Nature Neuroscience. 2020;23: 583–593. doi:10.1038/s41593-020-0603-0

12. Kichaev G, Yang W-Y, Lindstrom S, Hormozdiari F, Eskin E, Price AL, et al. Integrating Functional Data to Prioritize Causal Variants in Statistical Fine-Mapping Studies. PLOS Genetics. 2014;10: e1004722. doi:10.1371/journal.pgen.1004722

13. Li Y, Kellis M. Joint Bayesian inference of risk variants and tissue-specific epigenomic enrichments across multiple complex human diseases. 2016;44: 1–13. doi:10.1093/nar/gkw627

14. Ernst J, Kellis M. ChromHMM: automating chromatin-state discovery and characterization. Nature methods. 2012;9: 215–6. doi:10.1038/nmeth.1906

15. Wang D, Liu S, Warrell J, Won H, Shi X, Navarro FCP, et al. Comprehensive functional genomic resource and integrative model for the human brain. Science. 2018;362: eaat8464. doi:10.1126/science.aat8464

16. Myint L, Wang R, Boukas L, Hansen KD, Goff LA, Avramopoulos D. A screen of 1,049 schizophrenia and 30 Alzheimer’s-associated variants for regulatory potential. American Journal of Medical Genetics Part B: Neuropsychiatric Genetics. 2020;183: 61–73. doi:10.1002/ajmg.b.32761

17. Hoffman GE, Bendl J, Voloudakis G, Montgomery KS, Sloofman L, Wang Y-C, et al. CommonMind Consortium provides transcriptomic and epigenomic data for Schizophrenia and Bipolar Disorder. Sci Data. 2019;6: 180. doi:10.1038/s41597-019-0183-6

18. Won H, de la Torre-Ubieta L, Stein JL, Parikshak NN, Huang J, Opland CK, et al. Chromosome conformation elucidates regulatory relationships in developing human brain. Nature. 2016;538: 523–527. doi:10.1038/nature19847

19. Bernier R, Golzio C, Xiong B, Stessman HA, Coe BP, Penn O, et al. Disruptive CHD8 Mutations Define a Subtype of Autism Early in Development. Cell. 2014;158: 263–276. doi:10.1016/j.cell.2014.06.017

20. Stolerman ES, Smith B, Chaubey A, Jones JR. CHD8 intragenic deletion associated with autism spectrum disorder. European Journal of Medical Genetics. 2016;59: 189–194. doi:10.1016/j.ejmg.2016.02.010

21. O’Roak BJ, Vives L, Fu W, Egertson JD, Stanaway IB, Phelps IG, et al. Multiplex Targeted Sequencing Identifies Recurrently Mutated Genes in Autism Spectrum Disorders. Science. 2012;338: 1619–1622. doi:10.1126/science.1227764

22. Wilkinson B, Grepo N, Thompson BL, Kim J, Wang K, Evgrafov OV, et al. The autism-associated gene chromodomain helicase DNA-binding protein 8 (CHD8) regulates noncoding RNAs and autism-related genes. Transl Psychiatry. 2015;5: e568–e568. doi:10.1038/tp.2015.62

23. Sugathan A, Biagioli M, Golzio C, Erdin S, Blumenthal I, Manavalan P, et al. CHD8 regulates neurodevelopmental pathways associated with autism spectrum disorder in neural progenitors. Proc Natl Acad Sci U S A. 2014;111: E4468–4477. doi:10.1073/pnas.1405266111

24. Kogan CS, Turk J, Hagerman RJ, Cornish KM. Impact of the Fragile X mental retardation 1 (FMR1) gene premutation on neuropsychiatric functioning in adult males without fragile X-associated Tremor/Ataxia syndrome: a controlled study. Am J Med Genet B Neuropsychiatr Genet. 2008;147B: 859–872. doi:10.1002/ajmg.b.30685

25. Farzin F, Perry H, Hessl D, Loesch D, Cohen J, Bacalman S, et al. Autism spectrum disorders and attention-deficit/hyperactivity disorder in boys with the fragile X premutation. J Dev Behav Pediatr. 2006;27: S137–144. doi:10.1097/00004703-200604002-00012

26. Bourgeois JA, Cogswell JB, Hessl D, Zhang L, Ono MY, Tassone F, et al. Cognitive, anxiety and mood disorders in the fragile X-associated tremor/ataxia syndrome. General Hospital Psychiatry. 2007;29: 349–356. doi:10.1016/j.genhosppsych.2007.03.003

27. Clifton NE, Rees E, Holmans PA, Pardiñas AF, Harwood JC, Di Florio A, et al. Genetic association of FMRP targets with psychiatric disorders. Molecular Psychiatry. 2020; 1–14. doi:10.1038/s41380-020-00912-2

28. Folsom TD, Thuras PD, Fatemi SH. Protein expression of targets of the FMRP regulon is altered in brains of subjects with schizophrenia and mood disorders. Schizophr Res. 2015;165: 201–211. doi:10.1016/j.schres.2015.04.012

29. Kasap M, Rajani V, Rajani J, Dwyer DS. Surprising conservation of schizophrenia risk genes in lower organisms reflects their essential function and the evolution of genetic liability. Schizophr Res. 2018;202: 120–128. doi:10.1016/j.schres.2018.07.017

30. Pardiñas AF, Holmans P, Pocklington AJ, Escott-Price V, Ripke S, Carrera N, et al. Common schizophrenia alleles are enriched in mutation-intolerant genes and in regions under strong background selection. Nature Genetics. 2018;50: 381–389. doi:10.1038/s41588-018-0059-2

31. Song JHT, Lowe CB, Kingsley DM. Characterization of a Human-Specific Tandem Repeat Associated with Bipolar Disorder and Schizophrenia. Am J Hum Genet. 2018;103: 421–430. doi:10.1016/j.ajhg.2018.07.011

32. Xu K, Schadt EE, Pollard KS, Roussos P, Dudley JT. Genomic and Network Patterns of Schizophrenia Genetic Variation in Human Evolutionary Accelerated Regions. Mol Biol Evol. 2015;32: 1148–1160. doi:10.1093/molbev/msv031

33. Funk CC, Casella AM, Jung S, Richards MA, Rodriguez A, Shannon P, et al. Atlas of Transcription Factor Binding Sites from ENCODE DNase Hypersensitivity Data across 27 Tissue Types. Cell Rep. 2020;32: 108029. doi:10.1016/j.celrep.2020.108029

34. Nowakowski TJ, Bhaduri A, Pollen AA, Alvarado B, Mostajo-Radji MA, Di Lullo E, et al. Spatiotemporal gene expression trajectories reveal developmental hierarchies of the human cortex. Science (New York, NY). 2017;358: 1318–1323. doi:10.1126/science.aap8809

35. Bajic VB, Tan SL, Christoffels A, Schönbach C, Lipovich L, Yang L, et al. Mice and Men: Their Promoter Properties. Blake J, Hancock J, Pavan B, Stubbs L, PLoS Genetics EIC Wayne Frankel, editors. PLoS Genet. 2006;2: e54. doi:10.1371/journal.pgen.0020054

36. Lecellier CH, Wasserman WW, Mathelier A. Human enhancers harboring specific sequence composition, activity, and genome organization are linked to the immune response. Genetics. 2018;209: 1055–1071. doi:10.1534/genetics.118.301116

37. Vinogradov AE. Isochores and tissue-specificity. Nucleic Acids Research. 2003;31: 5212–5220. doi:10.1093/nar/gkg699

38. Cosgrove D, Whitton L, Fahey L, Broin PÓ, Donohoe G, Morris DW. Genes influenced by MEF2C contribute to neurodevelopmental disease via gene expression changes that affect multiple types of cortical excitatory neurons. Human Molecular Genetics. 2020. doi:10.1093/hmg/ddaa213

39. Mitchell AC, Javidfar B, Pothula V, Ibi D, Shen EY, Peter CJ, et al. MEF2C transcription factor is associated with the genetic and epigenetic risk architecture of schizophrenia and improves cognition in mice. Molecular Psychiatry. 2018;23: 123–132. doi:10.1038/mp.2016.254

40. Rocha H, Sampaio M, Rocha R, Fernandes S, Leão M. MEF2C haploinsufficiency syndrome: Report of a new MEF2C mutation and review. Eur J Med Genet. 2016;59: 478–482. doi:10.1016/j.ejmg.2016.05.017

41. Bishop KM, Garel S, Nakagawa Y, Rubenstein JLR, O’Leary DDM. Emx1 and Emx2 cooperate to regulate cortical size, lamination, neuronal differentiation, development of cortical efferents, and thalamocortical pathfinding. The Journal of Comparative Neurology. 2003;457: 345–360. doi:10.1002/cne.10550

42. Shinozaki K, Yoshida M, Nakamura M, Aizawa S, Suda Y. Emx1 and Emx2 cooperate in initial phase of archipallium development. Mechanisms of Development. 2004;121: 475–489. doi:10.1016/j.mod.2004.03.013

43. Kobeissy FH, Hansen K, Neumann M, Fu S, Jin K, Liu J. Deciphering the Role of Emx1 in Neurogenesis: A Neuroproteomics Approach. Frontiers in molecular neuroscience. 2016;9: 98. doi:10.3389/fnmol.2016.00098

44. Gorski JA, Talley T, Qiu M, Puelles L, Rubenstein JLR, Jones KR. Cortical excitatory neurons and glia, but not GABAergic neurons, are produced in the Emx1-expressing lineage. The Journal of neurosciencelZ: the official journal of the Society for Neuroscience. 2002;22: 6309–14. doi:20026564

45. Boix CA, James BT, Park YP, Meuleman W, Kellis M. Regulatory genomic circuitry of human disease loci by integrative epigenomics. Nature. 2021;590: 300–307. doi:10.1038/s41586-020-03145-z

46. Amemiya HM, Kundaje A, Boyle AP. The ENCODE Blacklist: Identification of Problematic Regions of the Genome. Scientific Reports. 2019;9: 9354. doi:10.1038/s41598-019-45839-z

47. Quinlan AR, Hall IM. BEDTools: a flexible suite of utilities for comparing genomic features. Bioinformatics (Oxford, England). 2010;26: 841–2. doi:10.1093/bioinformatics/btq033

48. Doan RN, Bae BI, Cubelos B, Chang C, Hossain AA, Al-Saad S, et al. Mutations in Human Accelerated Regions Disrupt Cognition and Social Behavior. Cell. 2016;167: 341-354.e12. doi:10.1016/j.cell.2016.08.071

49. Vermunt MW, Tan SC, Castelijns B, Geeven G, Reinink P, De Bruijn E, et al. Epigenomic annotation of gene regulatory alterations during evolution of the primate brain. Nature Neuroscience. 2016;19: 494–503. doi:10.1038/nn.4229

50. Karczewski KJ, Francioli LC, Tiao G, Cummings BB, Alföldi J, Wang Q, et al. The mutational constraint spectrum quantified from variation in 141,456 humans. Nature. 2020;581: 434–443. doi:10.1038/s41586-020-2308-7

51. Wright CF, Fitzgerald TW, Jones WD, Clayton S, McRae JF, van Kogelenberg M, et al. Genetic diagnosis of developmental disorders in the DDD study: a scalable analysis of genome-wide research data. Lancet. 2015;385: 1305–1314. doi:10.1016/S0140-6736(14)61705-0

52. Satterstrom FK, Kosmicki JA, Wang J, Breen MS, De Rubeis S, An J-Y, et al. Large-Scale Exome Sequencing Study Implicates Both Developmental and Functional Changes in the Neurobiology of Autism. Cell. 2020;180: 568-584.e23. doi:10.1016/j.cell.2019.12.036

53. Palmer DS, Howrigan DP, Chapman SB, Adolfsson R, Bass N, Blackwood D, et al. Exome sequencing in bipolar disorder reveals shared risk gene AKAP11 with schizophrenia. medRxiv. 2021; 2021.03.09.21252930. doi:10.1101/2021.03.09.21252930

54. Stahl EA, Breen G, Forstner AJ, McQuillin A, Ripke S, Trubetskoy V, et al. Genome-wide association study identifies 30 loci associated with bipolar disorder. Nat Genet. 2019;51: 793–803. doi:10.1038/s41588-019-0397-8

55. Howard DM, Adams MJ, Clarke T-K, Hafferty JD, Gibson J, Shirali M, et al. Genome-wide meta-analysis of depression identifies 102 independent variants and highlights the importance of the prefrontal brain regions. Nat Neurosci. 2019;22: 343–352. doi:10.1038/s41593-018-0326-7

56. Luciano M, Hagenaars SP, Davies G, Hill WD, Clarke T-K, Shirali M, et al. Association analysis in over 329,000 individuals identifies 116 independent variants influencing neuroticism. Nat Genet. 2018;50: 6–11. doi:10.1038/s41588-017-0013-8

57. Gandal MJ, Zhang P, Hadjimichael E, Walker RL, Chen C, Liu S, et al. Transcriptome-wide isoform-level dysregulation in ASD, schizophrenia, and bipolar disorder. Science. 2018;362. doi:10.1126/science.aat8127

58. Exome sequencing identifies rare coding variants in 10 genes which confer substantial risk for schizophrenia | medRxiv. [cited 13 Jun 2021]. Available: https://www.medrxiv.org/content/10.1101/2020.09.18.20192815v1

59. Genovese G, Fromer M, Stahl EA, Ruderfer DM, Chambert K, Landén M, et al. Increased burden of ultra-rare protein-altering variants among 4,877 individuals with schizophrenia. Nat Neurosci. 2016;19: 1433–1441. doi:10.1038/nn.4402

60. Pirooznia M, Wang T, Avramopoulos D, Valle D, Thomas G, Huganir RL, et al. SynaptomeDB: an ontology-based knowledgebase for synaptic genes. Bioinformatics. 2012;28: 897–899. doi:10.1093/bioinformatics/bts040

